# Generation and characterization of an ovine cell line derived from peripheral blood and its potential use to study livestock and zoonotic viral infections

**DOI:** 10.1101/2025.03.06.641825

**Authors:** S Moreno, C Sanjurjo-Rodríguez, S Rodríguez-Fernández, L Jiménez-Cabello, E Calvo-Pinilla, A Nogales, A Marín-López, J Ortego, A Brun, S Díaz-Prado, G Lorenzo

**Affiliations:** Centro de Investigación en Sanidad Animal CISA, INIA-CSIC. Valdeolmos (Madrid), Spain; Grupo de Investigación en Terapia Celular y Medicina Regenerativa, Instituto de Investigación Biomédica de A Coruña (INIBIC), Fundación Pública Gallega de Investigación Biomédica INIBIC, Complexo Hospitalario Universitario de A Coruña (CHUAC), Servizo Galego de Saúde (SERGAS). 15006 A Coruña, Spain; Grupo de Investigación en Terapia Celular y Medicina Regenerativa, Centro Interdisciplinar de Química y Biología (CICA), Departamento de Fisioterapia, Medicina y Ciencias Biomédicas, Facultad de Ciencias de la Salud, Universidade da Coruña (UDC), 15006 A Coruña, Spain; Centro de Investigación Biomédica en Red de Bioingeniería, Biomateriales y Nanomedicina (CIBER-BBN), 28029 Madrid, Spain; Department of Microbial Pathogenesis, Yale School of Medicine, New Haven, CT 06510, USA

**Keywords:** mesenchymal stromal cells, peripheral blood, cell-culture, viral infection, surface markers, differentiation, characterization, RT-qPCR

## Abstract

**Background:** Mesenchymal stromal cells (MSCs) are a population of undifferentiated non-hematopoietic fibroblast-like cells isolated from several tissues with multipotent differentiation capacity *in vitro*. This study focused on the establishment and characterization of a mesenchymal stromal cell line derived from ovine peripheral blood mononuclear cells (PBMCs).

**Methods:** MSC were isolated from ovine blood and used to develop a cell line. The characterization of the cell line to confirm its mesenchymal origin was carried out by different assays. First at all, cells were characterized by flow cytometry (FACS) using monoclonal antibodies specific for MSC and hematopoietic non-stromal cell surface antigens. Real-time quantitative PCR (RT-qPCR) was performed using gene primer pairs specific for hematopoietic and mesenchymal cell surface markers. In addition, we assayed the potential of the cell line to differentiate *in vitro* to other lineages such as osteoblasts and neurons and confirmed that by specific staining with Alizarin Red (AR) and RT-qPCR and immunofluorescence (IFA). Finally, we assayed the susceptibility of the cell line to various livestock, animal and human viruses.

**Results:** FACS and RT-qPCR analyses revealed that the ovine cell line expressed mesenchymal markers and was negative for hematopoietic markers assayed. MSC were able to differentiate to osteoblasts and neurons, these results were confirmed by quantitative analyses of AR staining and RT-qPCR and IFA assays. The cell line showed susceptibility to infection with all viruses assayed confirmed by IFA asssays.

**Conclusions:** In this work, we used isolated MSCs from peripheral blood to establish a new ovine cell line. The characterization and the osteogenic and neuronal differentiation carried out confirmed that the cell line had a mesenchymal stromal origin. In fact, the new established ovine cell line was permissive to all viruses studied in this work. The new sheep cell line could be a useful tool for isolating and characterizing viruses and studying virus-host interactions.

## 1. Introduction

Virus isolation is still used as the most specific method for diagnosis of viral infection to identify the viral agent on infected samples in veterinarian laboratories when the results of molecular testing or serology are unavailable or equivocal and it is also fundamental to generate a supply of virus for research purposes (1). Primary cells of bovine, chicken, ovine or porcine origin or continuous cell lines are used as a tool to isolate virus from biological samples or to produce large-scale antigen by diagnostic laboratories and veterinary pharmaceutical industries respectively (2). Considering primary cell cultures, they are difficult, expensive and time-consuming to produce, causing a delay in the diagnosis. When it comes to immortal cell lines the main disadvantage is that they are less susceptible to be infected, especially when its origin is different from the viral host range and this could cause an underestimation of possible positive cases, increasing the number of sick animals and the risk of an outbreak. These cell lines are also used to generate inactivated virus preparations used for vaccination purposes. Therefore, optimal viral growth is key to reduce costs related to vaccine production. The most common immortalized cell lines used to this purpose are Vero cells, derived from the kidney of an African green monkey; BHK-21 cells, derived from baby hamster kidney fibroblasts; and Madin-Darby Canine Kidney (MDCK) epithelial cells (3,4).

Mesenchymal stromal cells (MSCs) are a population of undifferentiated non-hematopoietic fibroblast-like cells isolated from several tissues (blood, adipose tissue, fetal liver, placenta, bone marrow, umbilical cord tissue) and from various animal species with multipotent differentiation capacity *in vitro* (5). Sheep and goats are commonly used as animal models for bone tissue engineering to test the potential of mesenchymal stromal cells for bone regeneration (6). MSCs have the ability to proliferate, adhere to plastic surfaces when cultured under standard conditions, express a certain panel of phenotypic markers and can differentiate into osteogenic, chondrogenic and adipogenic lineages when cultured in specific inducing media and other cell types like neuron-like cells, astrocytes or glial cells and corneal tissues (5,7–9). Due to their multi-differentiation potential and proliferative capabilities, they are potential candidates for cartilage and bone tissue engineering and are used in regenerative medicine to generate muscle, bones or as a tool to regenerate damaged tissues (10,11). In addition, immortalized MSCs lines can be helpful for the development of *in vitro* models for bone diseases and regeneration studies (12–14). Other studies have showed the therapeutic effects of MSCs being able to secrete active soluble substances to deliver immunomodulatory, antiapoptotic, angiogenic, and antioxidative effects (15).

Previous works have reported the isolation of MSCs from peripheral blood of different species including rabbits, mouse, rats, horses, dogs, humans and chickens (16–21). This procedure has several advantages like being less invasive and painful; it is easy and quick since it avoids other steps like enzymatic digestion or disaggregation to remove other contaminant tissues previous the MSCs purification; in addition to its easily growth under standard tissue culture conditions (22).

Other studies are focused on virus-MSCs interactions although there are limited information about the use of MSCs in infectious diseases (23–25). Particularly, there is scarce knowledge about the susceptibility to viral infection and antiviral properties, by upregulating their antiviral mechanisms, or effects on cell-based immune response (26).

In this work, we used isolated MSCs from peripheral blood to establish an ovine cell line to use it as a tool to isolate or produce virus from the same origin than the host. On the other hand, we studied the susceptibility of the new established ovine MSC line against a panel of relevant viruses causing infectious diseases.

## 2. Materials and Methods

### 2.1 Animals

All experiments with sheep were performed under biosafety level 3 (BSL-3) at INIA-CISA following EC guidelines [Directive 86/609) upon approval by the Ethical Review and Animal Care Committees of INIA and Comunidad de Madrid (authorization decision PROEX 192/17).

### 2.2 MSCs isolation and generation of MSCs culture

Peripheral blood mononuclear cells (PBMC) were obtained from a 9 months old female lamb (Castellana breed). Blood samples were collected from the jugular vein and immediately processed for the PBMC isolation. Briefly, 20 ml of blood were diluted in 1 volume of PBS (Phosphate-buffered saline buffer, Sigma) and carefully layered over Ficoll-Paque PLUS (GE Healthcare) in a 1:1 proportion and centrifuge at 400xg 30-40 min at 18-20°C. The mononuclear fraction was taken and rinsed twice in PBS by centrifugation for 5 min at 1600xg. Finally, cells were resuspended on DMEM (Dulbecco’s Modified Eagle’s Medium; Lonza, Spain) medium supplemented with 20% fetal bovine serum (FBS; Gibco, ThermoFisher Scientific, Spain), 1% penicillin/streptomycin (P/S; Gibco) and maintained at 37°C in 5% CO_2_.

Cells were initially seeded in 96 well tissue culture plates (Costar) on complete DMEM 20% FBS at 37° C with 5% CO_2_. Medium was replaced every 3-4 days removing non-adherent cells. When cells reached 80% confluence, they were detached using 0.25% Trypsin-EDTA (Sigma) and subcultured at a 1:2 ratio. Once the culture was expanded and the growth rate was stabilized, the percentage of FBS was reduced to 10%. The new cell line generated (called GeLo cell line or GeLo cells) were grown in chamber slides (Lab-Tek® Chamber Slide™ System – Nunc), fixed in iced methanol and stained with May-Grünwald Stain and observed under an inverted light microscope.

### 2.3 Cloning of GeLo cells

In order to get the culture homogeneity of the GeLo cell line, a method for cloning adherent mammalian cells was carried out (27). We obtained three different clones that were analyzed by FACS to check the presence and percentage of different mesenchymal markers present in their surface. Finally, both GeLo cell line and ovine bone marrow stromal cells (oBMSCs) were used for morphological, phenotypical and differentiation culture studies.

### 2.4 Culture of ovine bone marrow stromal cell line

Primary culture of oBMSCs was employed in this study as a control of MSCs cells. oBMSCs were cultured in complete 20%FBS-DMEM at 37° C in a humidified atmosphere with 5% CO_2_. The culture medium was then replaced every 3 days, when oBMSCs reached 80%, subculturing was performed for cell expansion using Trypsin-EDTA as mentioned above.

### 2.5 Phenotypic characterization using flow cytometry

GeLo cell line was trypsinized, washed and analyzed by FACS. A total of 2×10^5^ cells were transferred to FACs polypropylene tubes (NUNCTM, VWR International, Denmark). Optimal amounts of monoclonal antibodies (mAbs) were determined and added to each tube for 40 min at 4°C in darkness. The antibodies used are characteristic for markers associated with mesenchymal and hematopoietic lineages, some of them specific from human and others present cross reactivity with bovine and ovine and are listed in Table 1. A control tube for each of the chromogens used contained equivalent amounts of isotype standards. A minimum of 1×10^4^ cell events per assay was acquired using a FACsCalibur flow cytometer (BD Biosciences, Spain). Data were analyzed using Cell Quest software (BD Biosciences) and the results were expressed as mean of positive percentage.

**Table 1.** Monoclonal antibodies (mAb) used for flow cytometry.

| Antibody<sup>1 2</sup> | Clone | Specificity | Source |
| --- | --- | --- | --- |
| FITC Mouse IgG1 Kappa Isotype Control | MOPC-21 |  | BD Pharmingen |
| PE Mouse IgG1 Kappa Isotype Control | MOPC-21 |  | BD Pharmingen |
| PECy5 Mouse IgG1 Isotype Control | 1F8 |  | Abcam |
| LABELED PRIMARY ANTIBODIES | Clone | Specificity | Source |
| FITC Mouse Anti-Sheep CD44 | 25.32 | Homing cellular adhesion molecule (HCAM) | AbD Serotec |
| PE Mouse Anti-Human CD29 mAb | TS2/16 | $\beta 1$ Integrin | BioLegend |
| FITC Mouse Anti-Rat CD45 mAb | OX-33 | Leukocyte common antigen (LCA) | BD Pharmingen |
| PE Mouse Anti-Human CD73 mAb | AD2 | Esto-5'-nucleotidase | BD Pharmingen |
| PE-Cy5 Mouse Anti-Human CD90 mAb | 5E10 | Thy-1 membrane glycoprotein | BD Pharmingen |
| FITC Mouse Anti-Human CD105 mAb | SN6 | Endoglin, SH2 | AbD Serotec |
| PE Mouse anti-human CD166 mAb | 3A6 | Leukocyte-activated cell adhesion molecule (ALCAM) | Immunostep |

<sup>1</sup> Antibodies used for phenotypical characterization by flow cytometry. FITC=Fluorescein isothiocyanate;
PE= Phycoerythrin; PECy5= Phycoerythrin with cyanin-5.
<sup>2</sup> mAbs described in Khan et al (28).

### 2.6 Osteogenic differentiation of GeLo and oBMSCs

GeLo cell line and oBMSCs were differentiated towards osteoblast lineage. Differentiation experiments were carried out under normoxic conditions (i.e., a humidified atmosphere with 5% CO_2_ at 37°C). GeLo cell line and oBMSCs were detached using trypsin-EDTA, seeded at 2×10^4^ cells/well in a chamber slide (BD Falcon, France) for histology. Osteogenesis was induced by culture for 21 days in StemPro® Osteogenesis Differentiation Kit (Thermofisher Scientific), following the manufacturer’s instructions.

Culture media was changed every 2-3 days. Osteogenic differentiations were compared to a control consisting of oBMSCs cultured for the same period of time on 20%FBS-DMEM. Staining techniques were carried out to confirm the differentiation. For this purpose, cells were fixed in paraformaldehyde 4% and stained with Alizarin Red (AR, Sigma). For osteogenesis evaluation, differentiation was analyzed by AR staining, to assess the presence of calcium deposits.

### 2.7 Neuronal differentiation of GeLo cell line

GeLo cell line was chemically induced to become neuron-like cells *in vitro* using tetramethylpyrazine (TMP, Sigma) at 2.4 mg/ml in DMEM without FBS. After 24 hours of incubation, cells showed morphological changes similar to neurons and were visualized by a phase-contrast microscopy. Monolayer of differentiated cells were collected to isolate Poli A+ mRNA that was used to amplify specific markers (Table 2) expressed by neurons by RT-qPCR following manufacturer instructions (Qiagen).

**Table 2.**
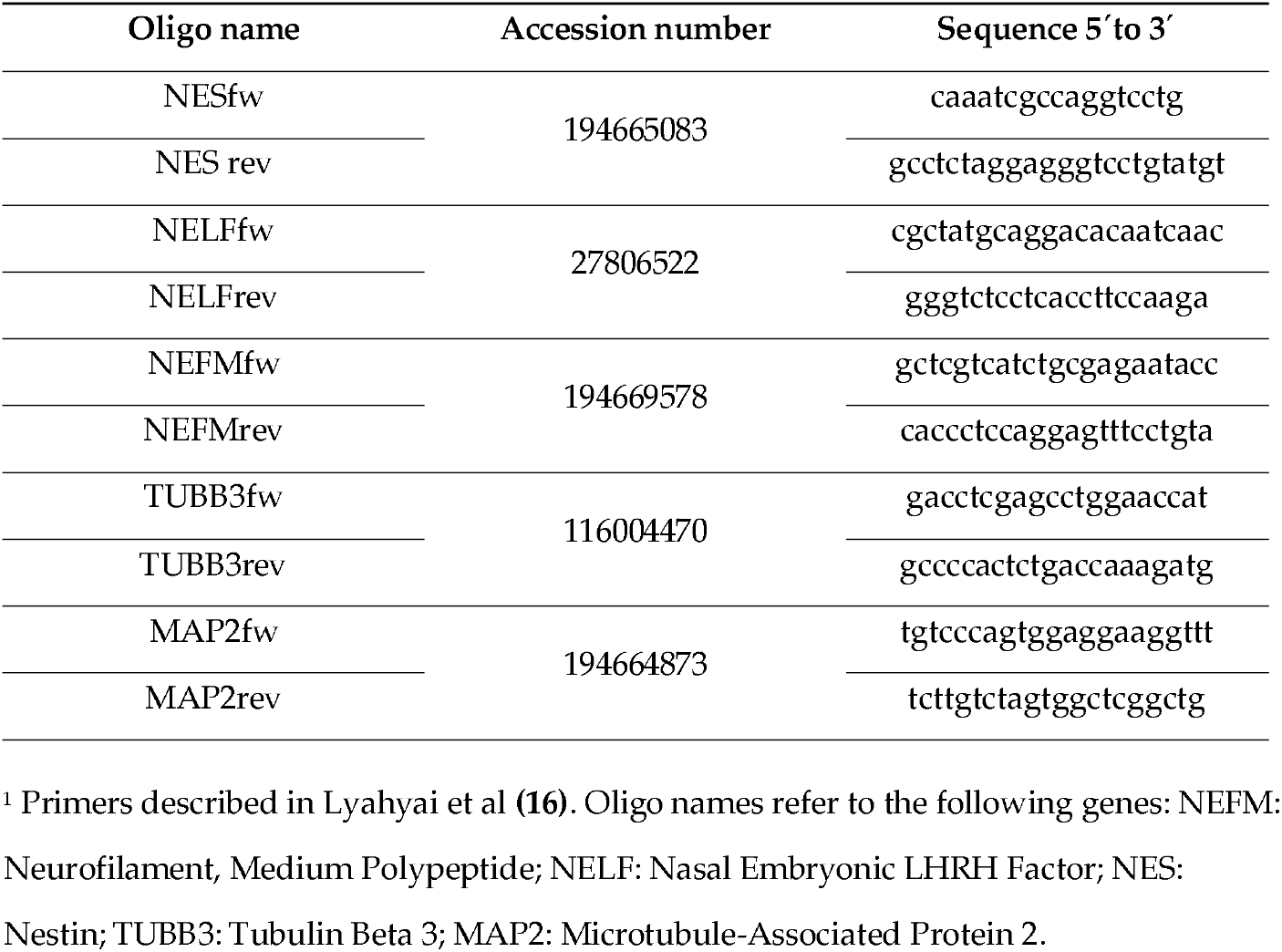
Differentiation neurogenic markers.

### 2.8 RNA extraction, reverse transcription and Real-time polymerase chain reaction (RTqPCR)

Poli A+ mRNA from GeLo cell line non and differentiated with TMPwere isolated from total RNA using Oligotex Direct mRNA minikit (Qiagen) and were reverse-transcribed into cDNA using Superscript IV reverse transcriptase (according to the manufacturer’s instructions, Invitrogen) in a thermocycler 2720 Thermal Cycler (Applied System). RTqPCR were performed using 1 μg of cDNA and SsoFast EvaGreen Supermix (Bio-Rad) in the Illumina·s Eco Real-Time PCR system (PCR-Max Limited, United Kingdom). The specific gene primer pairs sequences to mesenchymal, neuronal markers and those specific to hematopoietic markers employed are described in Table 2 and Table 3 respectively. Reference genes were used to normalize the expression. GAPDH (glyceraldehyde-3-phosphate dehydrogenase) and other genes as β-actin or Vimentin were used to analyze the expression levels. In the case of differentiated cells, the TUBB3 was used to normalize the expression levels. The thermal profile used consisted in heating (95°C/5s), followed by a stage cooling (60°C/5s) during 35 cycles and a melting stage of 95°C/5s, 65°C/5s and 95°C 5s. Data analysis was performed for triplicates and relative expression was calculated by comparing the levels of expression of the gene of interest against the levels of expression of an internal control gene, in this case vimentin. The formula used was ΔCT = CT(a target gene)-CT(a reference gene) (29).

**Table 3.**
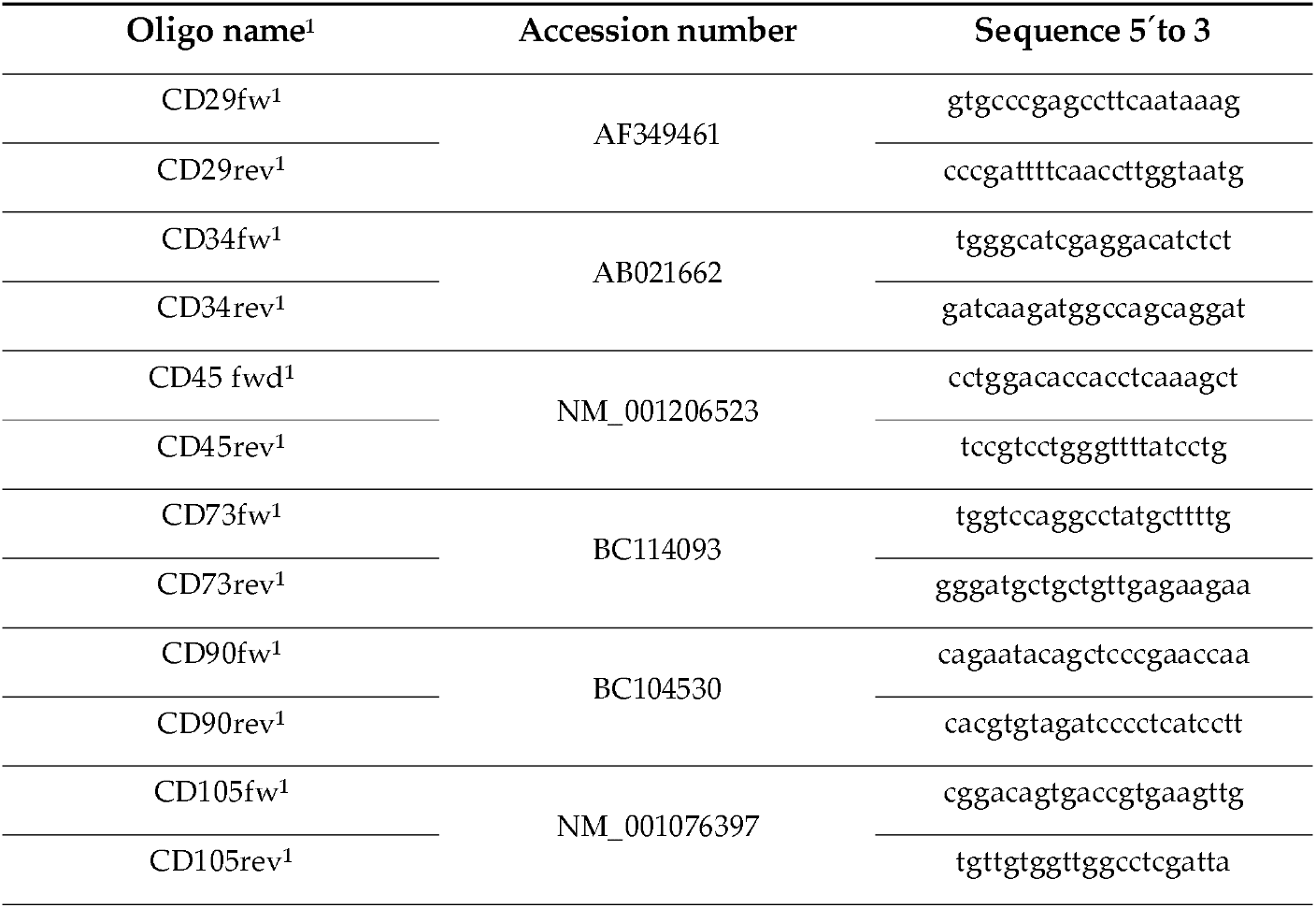
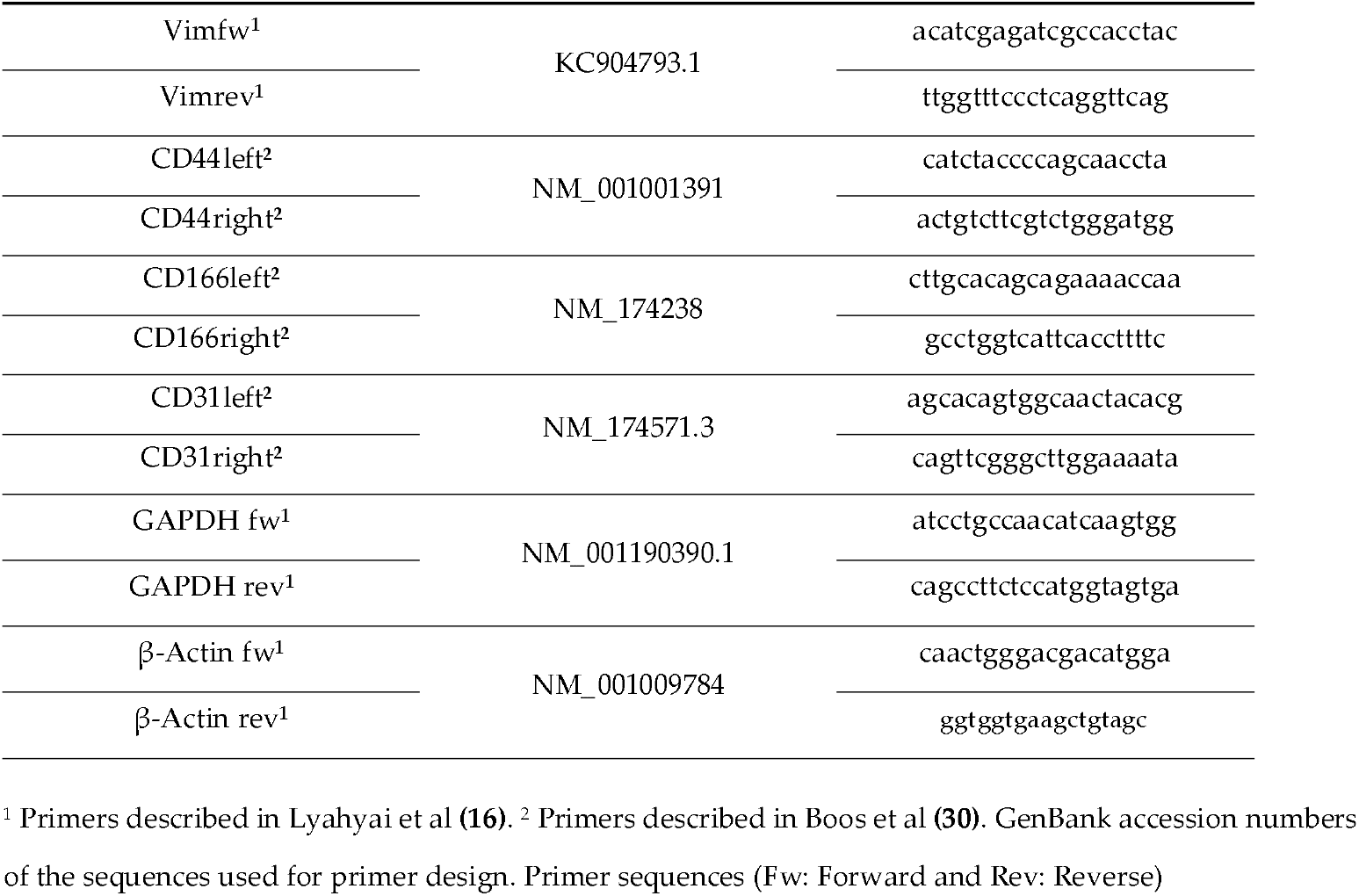
Primers used to amplify the PCR products by RT qPCR.

### 2.9 Virus infection assays and viral kinetics

GeLo cells were used to study their susceptibility to infection with several viruses causing disease in ruminants including Epizootic haemorrhagic disease virus (EHDV) responsible of a recent outbreak in Spain and Europe (31) and others affecting animals as well humans as Rift Valley fever virus (RVFV) or West Nile virus (WNV) (Table 4). Additionally, we assayed whether the viral vaccine vector Modified Vaccinia virus Ankara (MVA) (32) was able to infect the GeLo cell line with the purpose to adapt the viral vector to ovine cells and improve its potential as a vaccine vector in ruminants. For the infection assay, the GeLo cell line was infected to two multiplicity of infection (MOI) 0.1 and 0.01 and incubated one hour at 37°C, 5% CO_2_ and 95% humidity atmosphere in a cell incubator. After 1 h of viral adsorption, the cells were supplemented with fresh medium containing 2% FBS and then were maintained at 37ºC. Then the cells were observed every 24 hours under a phase-contrast inverted microscope to detect cytopathic effect (CPE). In the case we observed CPE, the supernatant was collected and titrated, to this end Vero cells were seeded in 24-well plates and inoculated at ten-fold serial dilutions in DMEM 2% FBS. After 1 h incubation at 37°C, the inoculum was removed and cells were overlaid with semi-solid medium (Eagle’s Minimum Essential Medium (EMEM)-1% carboxymethylcellulose) and further incubated for 3 days at 37°C to allow plaque development. Afterwards, the semi-solid overlays were poured off with water and the monolayer was fixed with 10% formaldehyde and stained with 2% crystal violet. Viral titers were calculated from the plaque numbers obtained for each dilution.

**Table 4.** Virus employed in this work and their host range.

| Virus | Strain | host |
| --- | --- | --- |
| Rift Valley fever virus (RVFV) | MP12 | Domestic and wild animals;<br>humans |
| Schmallenberg virus (SBV) | BH | ruminants |
| Toscana virus (TOSV) | ISS Phl.3 | humans |
| Akabane virus (AKV) | JaGAR-39 | ruminants |
| West Nile virus (WNV) | NY99 | humans, equines and some<br>birds |
| Peste du Petit Ruminants virus (PPRV) | Morocco | ruminants |
| Foot and Mouth disease virus (FMDV) | O | cloven-hoofed animals<br>(cows, pigs, sheep, goats,<br>and deer) |
| African horse sickness virus (AHSV) | AHSV-4 | all species of Equidae |
| Modified Vaccinia virus Ankara (MVA) | Ankara | Adapted to growth in avian<br>cells |
| Epizootic hemorrhagic disease virus<br>(EHDV) | EHDV-8 | wild and domestic<br>ruminants |
| Influenza A Virus (AIV) | A/PR/8/1934 H1N1 | Avian and mammal hosts |
| Blue Tongue virus (BTV) | BTV-4 | ruminants |

Once, the viral infection was confirmed with all viruses from Table 4, a growth kinetics assay was carried out to compare the virus production in the new established cell line with common cell lines used to this purpose like Vero cells or BHK-21 cells. For this aim, cells seeded in 24- well plates at a density of 4 × 10^5^ cells/mL in DMEM 2% the day before, were infected with the different viruses mentioned above at a MOI of 0.1 or 0.5 with the exception of WNV strain New York with a MOI of 0.01 due to its fast growth. A different assay was employed with Influenza A virus as described (33). Supernatants were collected at 24, 48 and 72 hours post infection, except for FMDV since it grows faster than the others, to determine the infection kinetics of each virus in two cell lines, their traditional growth cell line and GeLo. Finally, the titer of each supernatant was assayed in Vero cells by plaque-assay as described previously.

### 2.10 Immunofluorescence and confocal microscopy

GeLo cells were infected at MOI of 1 in glass slides (15mm Diameter, Fisher Scientific). After 24h, GeLo cells were fixed with 10% paraformaldehyde for 15 minutes and washed with PBS and blocked with 20% FBS in PBS (blocking and dilution buffer) for 1h at RT. Next, cells were incubated with anti-sera specific for each virus obtained from experimental infected mice, rabbit or sheep and natural infected sheep in the case of EHDV. After two washes with dilution buffer, cells were incubated in the same buffer with a secondary antibody conjugated with Alexa-488 specific for mouse or sheep Immunoglobulins G (IgG) for 1 h at 37^°^C. Slides samples were examined by confocal microscopy using a microscope Zeiss LSM880 confocal laser microscope (Gmbn).

### 2.11 Cytokine Analysis in GeLo cell cultures

GeLo cell cultures were treated with phytohemagglutinin (PHA, as positive mitogen control) or infected with diferent DNA or RNA viruses for 24 h. Then, supernatants were collected and used in the assays. Cytokine levels were analyzed using a multiplex fluorescent bead immunoassay (MILLIPLEX Mouse Cytokine kit, Burlington, MA, USA) for quantitative detection of ovine cytokines (IL-1β, IL-4, IL-6 and IL-8) following the manufacturer’s instructions. Samples were analyzed with a MAGPIX system (LuminexCorporation, Austin, TX, USA) and data were collected using xPonent version 3.1 software. The cytokine concentrations (pg/ml) were calculated using standard curves. Values that fell below the level of detection of the assay were assigned the lowest detectable concentration.

## 3. Results

### 3.1 MSCs isolation, culture and morphology

MSCs cells were selected by plastic adherence as described previously (34). The GeLo cell line cultured *in vitro* observed in a phase-contrast inverted microscope (Nikon TMS) presented a homogenous population with a typical spindle-shaped morphology with polygonal shapes. Cells showed an irregular cytoplasm with numerous branches and oval nuclei with prominent nucleoli and fine chromatin characteristic of the MSCs (Figure 1a).

**Figure 1.**
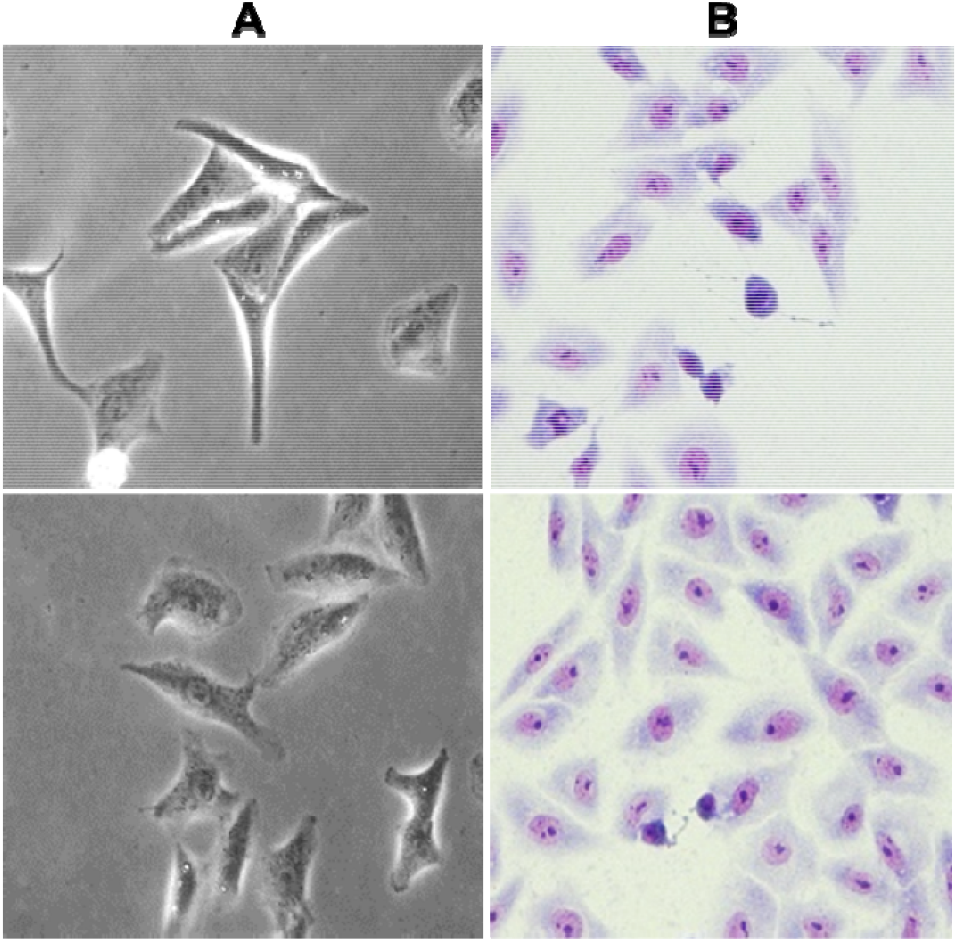
A. Morphological phenotype of established GeLo cell line derived from peripheral blood in culture. Images were taken under an inverted light microscope at 20x magnification. B. May-Grunwald Giemsa staining of cells showed the cytoplasm stained in blue with the nucleus stained in pink with two centrioles.

To analyze more in detail their morphology, GeLo cells were stained with May-Grünwald Giemsa Stain. This stain shows the nucleus in pink with two centrioles, the nuclear DNA and chromatin in purple and the cytoplasm stained in blue (Figure 1b). The GeLo cell line presented high cellular activity with numerous cells in cellular division (mitosis).

### 3.2 Phenotypic characterization

Immunophenotypic analysis was carried out to test the expression of mesenchymal and hematopoietic markers in the newly established GeLo cell line. Cells were incubated with a panel of antibodies characteristic of MSCs (CD29, CD44, CD73, CD90, CD105 and CD166) and hematopoietic cells (CD45). These antiboides were specific of sheep or other species such as human, mouse or rat but cross-reactive with ovine cells (Table 1).

By FACS analysis, we observed that GeLo cells were positive for MSC markers like CD29, CD44 and CD166, and negative for CD45, CD73, CD90 and CD105 (Figure 2). At the same time, the percentage of cells positive to different MSCs markers after the cloning of the GeLo cell line was also analyzed. GeLo cell line showed values between 66 to 97% after cloning with the different surface markers employed in comparison with those obtained before 26-28% (supplementary figure 1) confirming the success of the cloning assay (Figure 2).

**Figure 2.**
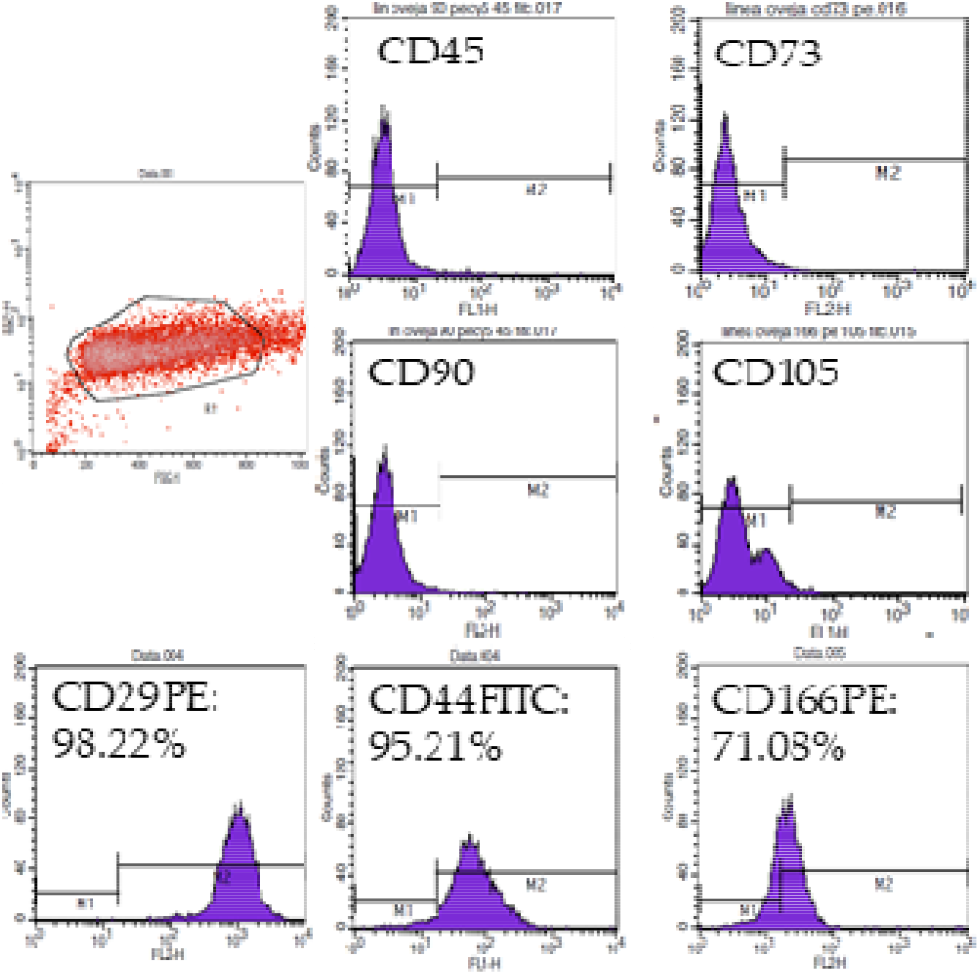
Phenotypic characterization by flow cytometry of GeLo cell line (after cloning) for markers characteristic of MSCs (CD29, CD44, CD73, CD90, CD105 and CD166) and hematopoietic cells (CD45). GeLo cell line was positive for CD29, CD44 and CD166 and negative for CD45, CD73, CD90 and CD105.

### 3.3 Osteogenic differentiation

To study the potential of the GeLo cell line to differentiate to other cell lineages, we first analyzed its capacity to differentiate to osteoblasts. To that end, we incubated GeLo cells with StemPro® Osteogenesis Differentiation Kit and evaluated the newly formed calcium deposits by staining with Alizarin Red (AR) at 28 days post-differentiation. GeLo cell line presented nodule-like aggregations stained in red when they were maintained with osteogenic medium in comparison with those cells with DMEM 20% FBS or oBMSCs used as a control grown in both media (Figure 3A) where a faint stain or no deposits were observed. These results were confirmed by quantitative analyses of the AR staining using ImageJ 1.48v (National Institutes of Health, Bethesda, USA) (Figure 3B).

**Figure 3.**
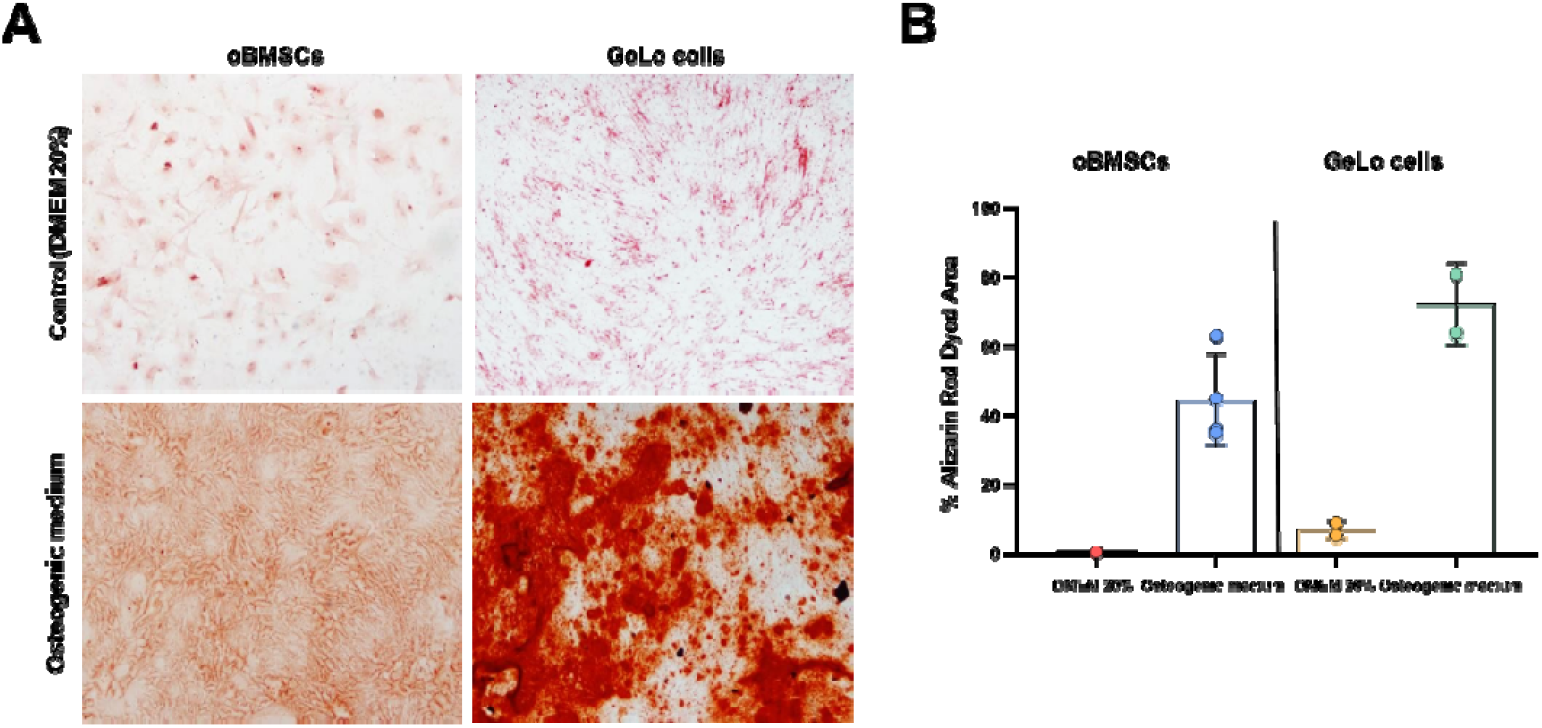
Quantitative analyses of the Alizarin Red staining were measured using ImageJ 1.48v (National Institutes of Health, Bethesda, USA). After the colour substraction of non-stained areas, the percentage of metachromatic areas was measured and expressed as mean±standard error.

### 3.4 Neuronal differentiation and analysis of the expression of gene markers

We also studied the potential of the GeLo cell line to differentiate to neuron-like cells. After 24 hours in culture with TMP, cells presented a drastic change in their morphology. The neuron-like phenotype was confirmed by: (i) morphological analysis (Figure 4A); (ii) immunofluorescence with a neuronal-specific protein/marker namely β-tubulin III (a microtubule structural marker expressed earliest phases of neuronal differentiation), (Figure 4B), (iii) expression of different neuronal markers (βTUBIII, NELF, NES, MAP2) by q RT PCR (Figure 4D).

**Figure 4.**
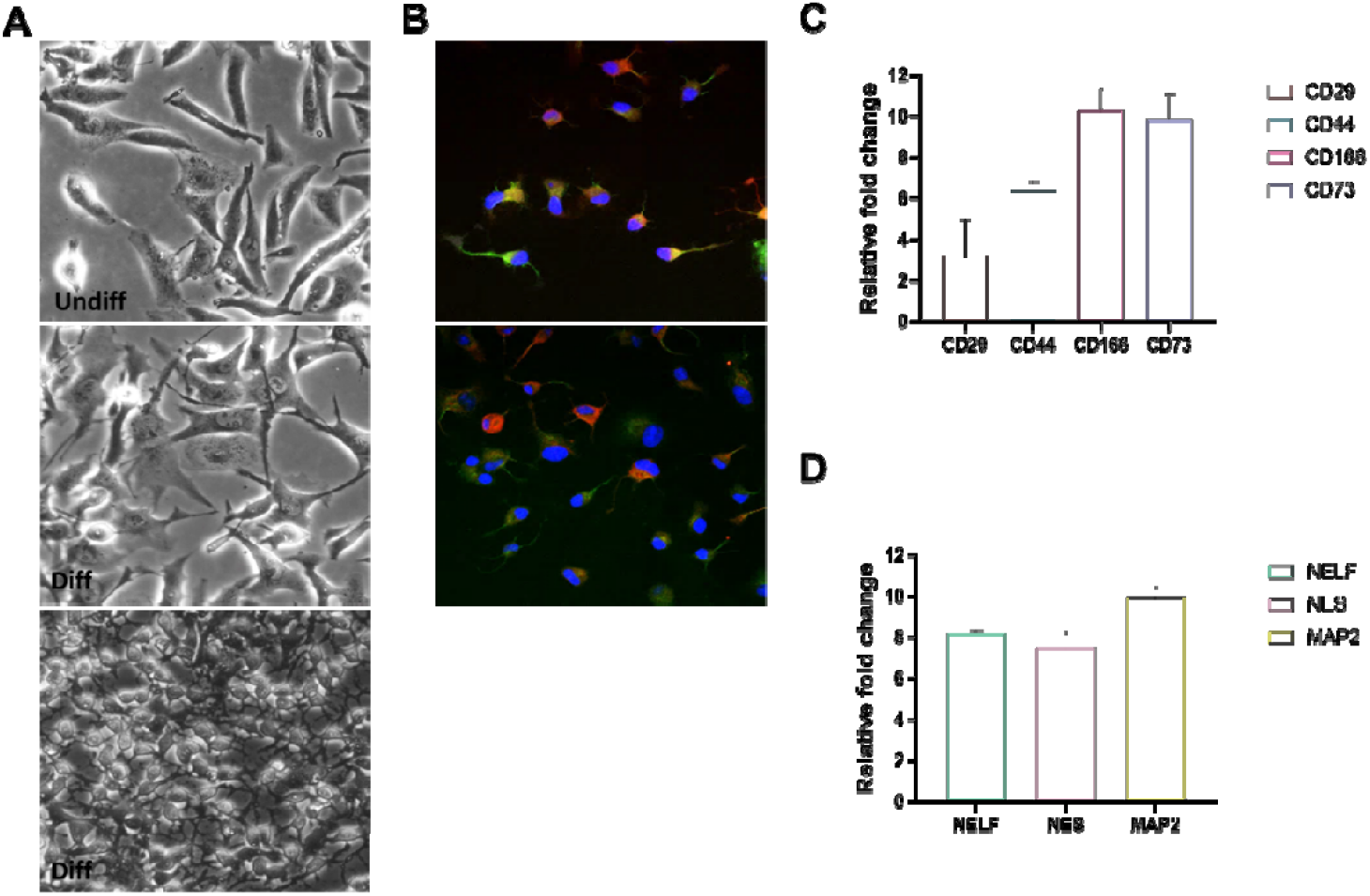
**A**. GeLo cell line differentiation in the presence of TMP. Cells show a neuro-like phenotype. **B**. Cells were marked using an anti-tubulin β-3 (Red) and anti-vimentin (Green). 63x. **C**. Analysis by RT- qPCR of MSCs surface markers expression, the ovine cell line expressed efficiently CD29, CD44, CD166 and CD73, all of them surface markers of MSCs. **D**. Analysis by RT qPCR of neuronal markers expression during neuronal differentiation, GeLo cell line expressed efficiently specific markers of neurons (NELF, NES, MAP2 and TUBB3) after differentiation in culture in the presence of neurobasal medium supplemented with B-27 and GlutaMAX-I. Bars represent the mean value and error bars represent the standard deviation.

GeLo cell line showed a reduction in size with a rounded bright nucleus with fine and long multipolar extensions and branching end brands (Figure 4A). To confirm the neuronal differentiation of GeLo cells, an immunofluorescence assay was performed to identify the expression of neural specific markers using specific mAbs to detect β-tubulin III (a marker of mature neurons) and vimentin (type III intermediate filament protein expressed in mesenchymal cells). Many of the differentiated GeLo cells expressed the neuronal marker β- tubulin III (red) or vimentin (green) and some of them co-expressed both proteins (merged in yellow) (Figure 4B). These results confirmed the potential of the GeLo cell line to differentiate into neuron-like cells *in vitro* in the presence of TMP.

Thereafter, we analyzed the expression of gene markers before and after differentiation of GeLo cells by RT-qPCR of each target gene (Table 2 and 3). Vimentin was used as the reference gene based on the results obtained by RT-PCR with several housekeeping genes (HK). This gene showed a stable expression compared with the variability or no expression of HK genes (GAPDH or β- actin) after several rounds. Results obtained with mesenchymal primers showed that GeLo cell line expressed efficiently CD29, CD44, CD73 and CD166 being the CD29 the most expressed and CD166 the less one confirming its mesenchymal origin/lineage (Figure 4C). In addition, several markers specific of neuronal cells (βTUBIII, NELF, NES, MAP2) were used by RT-PCR in the GeLo cell line and after TMP differentiation. Differentiated GeLo cell line expressed the neuronal markers NELF, NES and MAP2 with expression levels very similar (figure 4D) in contrast with undifferenciated GeLo cells that expressed none of those markers before. These results are consistent with those obtained by FACS.

### 3.5 Detection of viral antigens in GeLo cells by immunofluorescence

We wondered whether GeLo cells were susceptible to infection with a different viruses, including zoonotic or whose host range is restricted to ruminants (see Table 4). To that end, GeLo cells were infected with RVFV, AkV, BTV-4, MVA, AHSV-4, EHDV-8, TosV, FMDV, SBV, PPRV, WNV and IAV and CPE was monitored. To confirm the presence of the antiviral antigens, indirect immunofluorescence assays (IFA) were performed to detect viral antigens in infected cells by means of the use of specific sera or monoclonal antibodies against each of the viruses tested. Results confirmed the efficient expression of viral antigens in the cytoplasm of the GeLo cell line. Protein expression was detectable at 24 hpi. (figure 5). Altogether, these results confirmed that the GeLo cell line is susceptible to viral infection.

**Figure 5.**
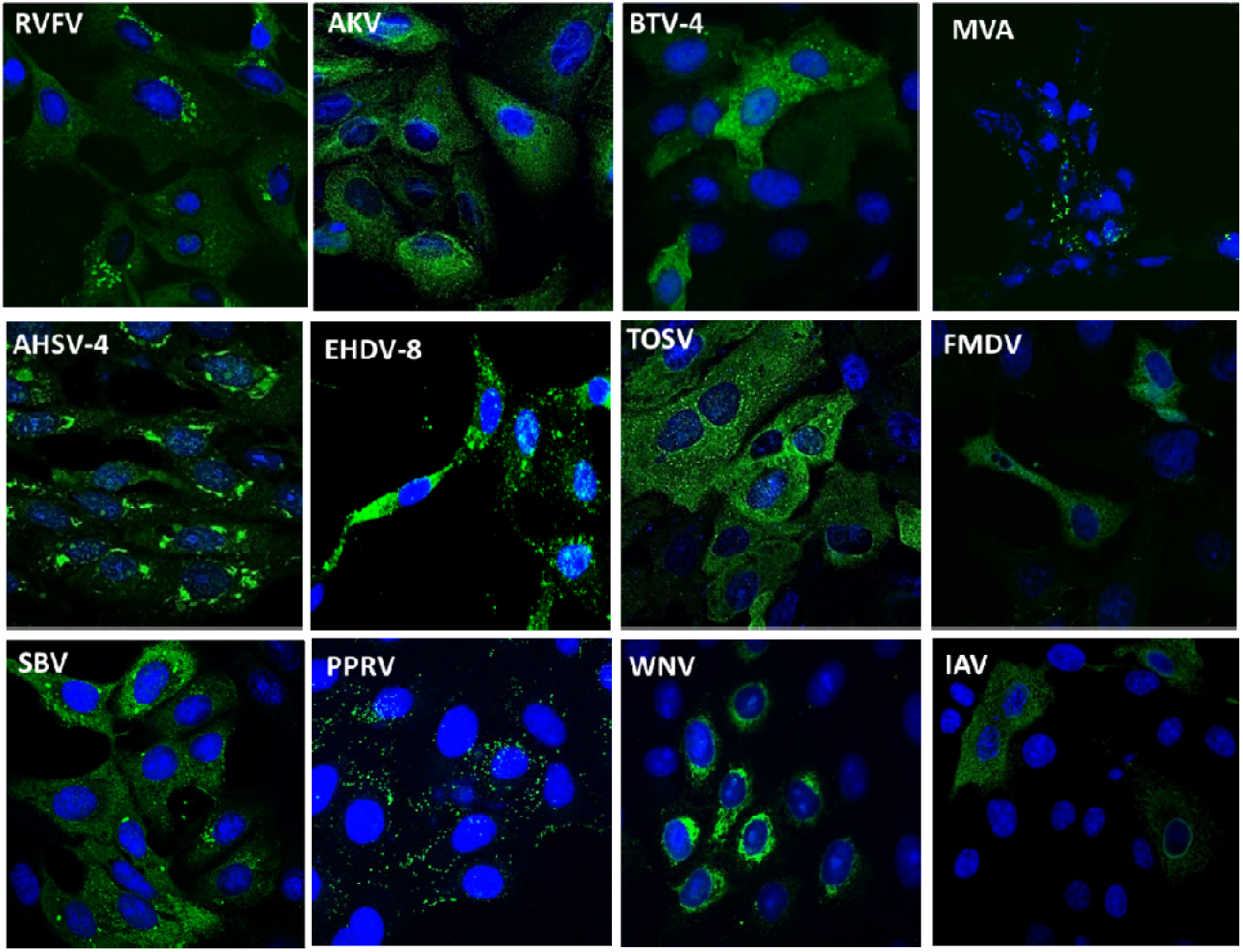
Indirect immunofluorescence in infected GeLo cells. Cells were infected with different viruses with a MOI of 1 and fixed 24 hpi. Green fluorescence (Alexa 488) shows viral infection using positive sera or monoclonal antibodies against the virus antigens. DAPI staining was used to show cell nuclei. Images were taken by confocal microscopy (63x magnification).

### 3.6 Susceptibility of the GeLo cell line to viral infection

To confirm infection and the ability of these viruses to replicate in GeLo cells the growth kinetics of the different viruses were monitored. GeLo cells were susceptible to infection with all these viruses after 48h from the first passage (data not shown), with the exception of Schmallenberg virus where it was necessary to carry out one blind passage to obtain infectious virus in the following passage. Syncytia formation during PPRV infection was observed after 4 days of culture in GeLo cells (Supplementary figure 2) confirming the success of viral infection. The growth kinetics assay was also performed to compare the virus production in the GeLo cell line with other cell lines commonly used to produce virus as Vero cells and BHK-21 cells. Results showed similar curves in most cases being the GeLo cells production lower during the first 24 h and obtaining similar values to Vero or BHK-21 cells in the next time points, apart from MVA where values are significantly lower in GeLo cells at all time points. The viral titre at 72 hpi is almost identical to the original viral input (Figure 6). Despite productive infection occurring in GeLo cells, FMDV and SBV virus growth was slowed down in GeLo cells compared to that observed in BHK-21 or Vero cells, respectively (Fig. 6). The variation observed after 72h with RVFV infection was associated with the state of the infected monolayer after three days in culture.

**Figure 6.**
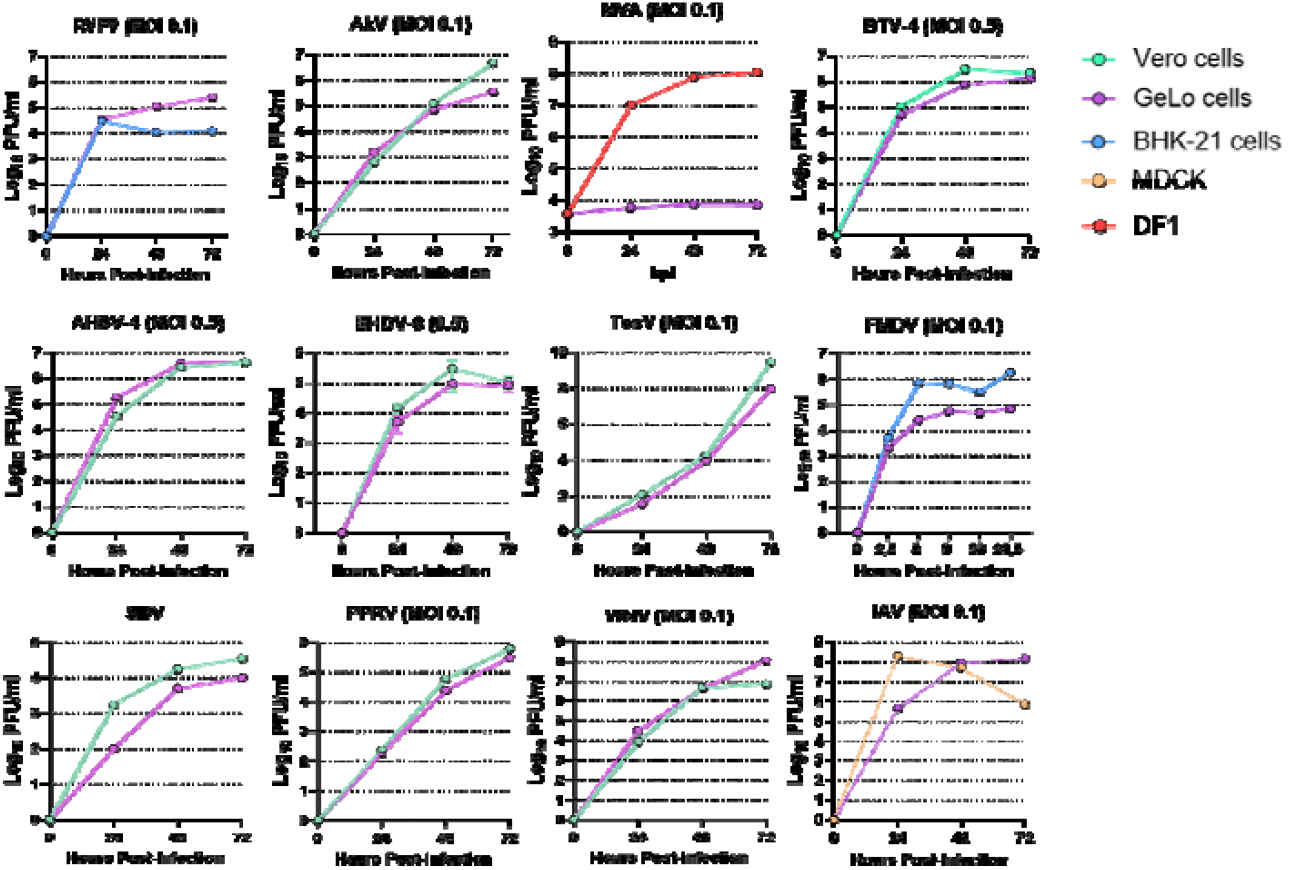
Viral kinetics of different viruses were performed in Vero, BHK-21 or MDCK cells and GeLo cell line at three different time points (24, 48 and 72 hpi, hours post infection). MOI is indicated in brackets. Results showed that GeLo cell line was susceptible to infection with all viruses employed (Table 4).

### 3.7 Cytokine Analysis in GeLo cell cultures

As mentioned above, MSCs have anti-inflamatory and immunomodulatory potential since they can regulate the secretion of proinflamatory and antiinflamatory cytokines such as IL-1B or IL-6 respectively (A). During the beginning of infections MSCs are activated by viral antigens eliciting a strong immune response by producing proinflammatory cytokines which enhance the antiviral properties of immune cells (35). Thus, we investigated whether GeLo cells are capable of eliciting immunomodulatory responses by secreting cytokines, such as IL-1B, IL-4, IL-6 and IL-8, with antiviral properties. To this end, we performed infection assays with RNA or DNA viruses in GeLo cells. The results (Figure 7) showed that supernatants from GeLo cells, infected DNA viruses, had detectable levels of IL-1β, IL-4, IL-6 and IL-8. Protein levels of IL-1β were not detectable in supernatants from GeLo cells infected with RNA viruses; Levels of IL-4, IL-6 and IL-8 were detectable, similar for IL-6 and IL-8, and lower for IL-4.

**Figure 7.**
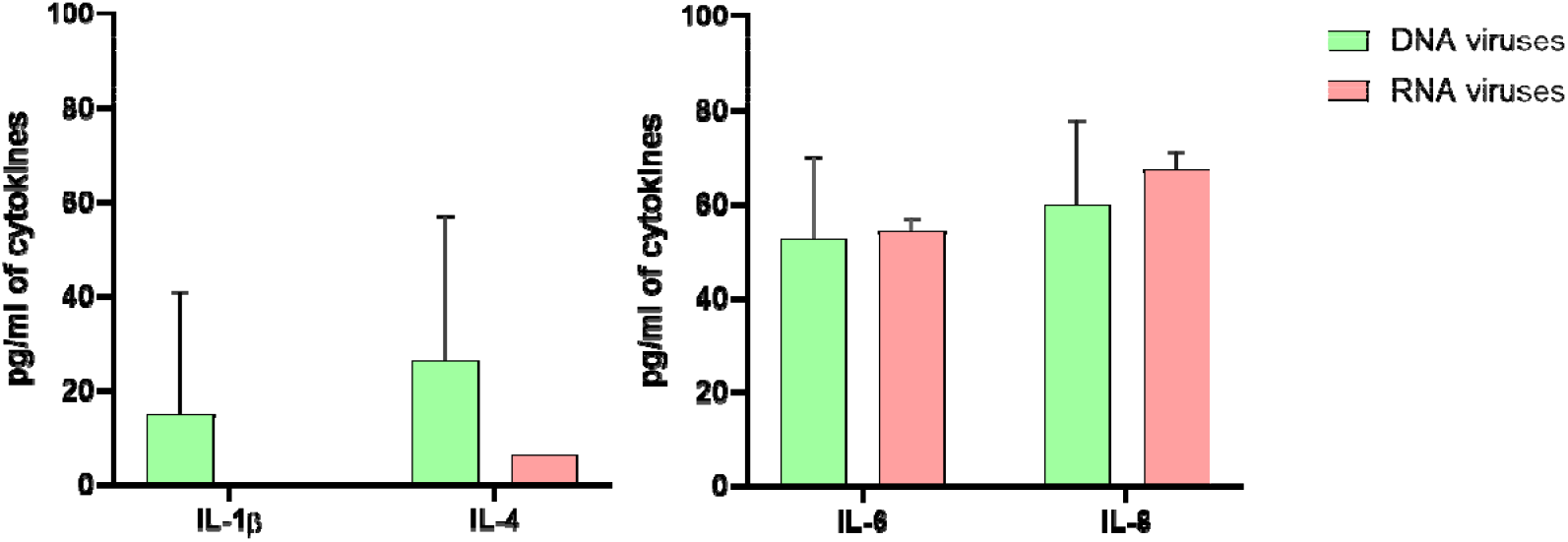
Cytokine secretion levels by GeLo cells after stimulation with DNA or RNA viruses. After 24 h, supernatants were collected and cytokines determined by Luminex assay (IL-1β, IL-4, IL-6 and IL-8). Values of cytokines from media-stimulated samples were subtracted from values of virus stimulated samples.

## 4. Discussion

MSCs can be isolated from different tissues such as bone marrow, adipose tissue, umbilical cord or peripheral blood from different species (5,6,8). Human MSCs are the best characterized, and the International Society for Cellular Therapy has proposed minimal criteria for defining this type of cells (7). First, MSCs should adhere to plastic; second, MSCs should express surface markers of mesenchymal stromal cells CD105, CD73 and CD90 and should not express CD45, CD34, CD14 or CD11b, CD79a or CD19 and HLA-DR; and finally, MSCs should be able to differentiate into osteoblasts, adipocytes and chondroblasts *in vitro* (9–11). The GeLo cell line was characterized following the same criteria as human MSCs.

In this study, fibroblast-like cells were isolated from peripheral blood by adherence to plastic. Several rounds of washings were carried out to remove non-adherent cells while they were maintained in 20%FBS DMEM. Once the culture was stabilized, these cells were expanded, and different assays were performed to carry out their characterization. The expression of mesenchymal markers and absence of hematopoietic markers was analyzed and confirmed by flow cytometry and RT-qPCR. The ability of differentiation to osteoblasts or neurons was carried out by *in vitro* culture, RT-qPCR and immunofluorescence. On the other hand, we studied the susceptibility of the newly established ovine mesenchymal stromal cell line against a panel of relevant viruses that cause infectious diseases.

The newly established ovine cell line (called GeLo cell line) showed certain cell type heterogeneity in culture. To solve this issue, adherent cells were successfully cloned by limiting dilution (27). Several clones were obtained and the immunophenotypic characterization was carried out by FACS. Despite the importance of sheep as a large animal model for many studies the characterization of ovine MSCs (oMSCs) is still limited. A major limitation is that there are no specific ovine reagents and there is little information about the surface markers present in oMSC. These cells have principally been isolated from human bone marrow, being well characterized by the surface markers specific expressed. We used a total of eight mAbs, some of them specific to human and others present cross reactivity with bovine and ovine (Table 1) (28). Notably, GeLo cell line expressed three MSCs markers (CD29, CD44 and CD166) and none of the hematopoietic cell markers. The negative results obtained for several mesenchymal surface markers, such as CD45, CD73, CD90 and CD105, could be due to the absence of reactivity of these mAbs in sheep since many of these antibodies are specifically designed or optimized for humans. Even the isolation and analysis methods could influence the detection of these markers, for example the culture medium or the available oxygen could modify the percentage and type of cells obtained (10,15,16,28,36) and a less sensible analysis methods might not detect markers with a low level of expression. Another possibility is the absence of these markers in oMSCs from blood since it is known that the expression of these markers could vary depending on the tissue of origin. Furthermore, MSCs are not homogenous, in a population, not all cells express uniformly CD73, CD90 and CD105; it is possible that a subpopulation of MSCs have a low expression of these markers or not expression at all. On the other hand, controversy exists among the results because depending of the specie where MSCs have been isolated could express or not the markers described above. There are studies where they identify markers in other species that are not characteristic in hMSCs (37,38); there might be specific proteins that do these functions in sheep. It is clear that there are several factors that might cause the lack of expression of CD73, CD90 y CD105 and it does not necessarily mean that these cells are not MSCs, as long as a functional evaluation, as the one describe in this manuscript, and a neuronal and osteogenic differentiation is done. In our experience, we were not able to detect CD73 by FACS but we obtained CD73 transcripts by RT qPCR confirming their expression. A possible explanation for CD73 is that their clones displayed variable degrees of binding affinity and in our assays the mAb used did not work. Another variable is the method used to detach the cells from the tissue culture plastic that can digest some cell-surface markers, leading to their loss or reduced detection during FACS (28,36,39).

We also analyzed the expression of mesenchymal and hematopoietic markers by RT-qPCR in GeLo cells. We used a list of ten pairs of primers specific of ovine or bovine surface markers. There was no amplification of the hematopoietic markers CD45, CD31 or CD34 confirming the results obtained by FACS. We confirmed the expression of CD29, CD44 and CD166 and also detected transcripts of CD73 by RT qPCR in the GeLo cell line. Due to the poor availability of ovine reagents, RT-qPCR could be an interesting alternative to characterize gene expression of surface markers in MSCs (22,40).

The second goal of this work was to carry out the osteogenic and neuronal differentiation to demonstrate the capability of the GeLo cell line to differentiate in other lineages typical of MSCs. The differentiation into osteoblasts was demonstrated using AR (Figure 3a). Nodule-like aggregations stained in red appeared in the osteogenic media on the 21st day of culture, indicating that these cultures were mineralized at a late stage. These results are compatible with previous published studies (10,16,28).

We also carry out neuronal differentiation in the presence of TMP. In only 24h GeLo cells morphology was drastically modified. We confirmed the process by RT-qPCR using specific primers of neuronal markers. We detected expression profiles of neurogenic markers (MAP2, NELF, NES and TUBB3) that are consistent with neural differentiation. TUBB3 was used to normalize the results obtained due to the variability obtained with HK genes. We confirmed the results obtained by RT-qPCR by immunofluorescence. GeLo cells differentiated to neurons were stained with Vimentin and TUBB3, we observed single and double stained cells in a high percentage of them. These results confirmed the potential of the GeLo cell line to differentiate into neuron-like cells *in vitro* in the presence of TMP (9).

Finally, we studied whether the GeLo cell line was susceptible to infection with a panel of relevant DNA and RNA viruses causing infectious diseases in animals or/and humans. Interestingly, results obtained by viral inoculation showed that GeLo cells were susceptible to infection with all viruses studied in this work. The capacity of these virus to infect GeLo cells was determined by immunofluorescence for the detection of viral antigens with specific sera or mAbs (figure 5). And the success of the viral infection was confirmed by a growth kinectics assay (figure 6). Previous studies have demonstrated that MSCs are susceptible to infection by a wide variety of RNA and DNA viruses both *in vitro* and *in vivo*. The capacity of viruses to enter and infect host MSCs may also be host species dependent (26). MSCs have many functional surface receptors, which potentially could enable viral entry. MSCs surface receptor expression and viral tropism may in part clarify MSCs susceptibility to viral infection. MSCs are highly permissive to infection by Herpesviruses, including Herpes Simplex-1 (HSV-1), Cytomegalovirus (CMV) and Varicella Zoster virus (VZV) (41). As mentioned before, some studies have demonstrated the therapeutic effects of MSCs being able to secrete active soluble substances to induce immunomodulatory responses (42,43). MSCs that are activated by viral antigens elicit strong immune responses by production of proinflammatory cytokines which enhance the antiviral properties of immune cells at early stages of infection (35). Therefore, we decided to assess whether GeLo cells were capable of having immunomodulatory responses by secreting cytokines with antiviral properties, such as IL-1B, IL-4, IL-6 and IL-8. For that purpose, we carried out infection assays with RNA or DNA viruses in GeLo cells.

For Il-6 and IL-8, results showed that GeLo cells display a differential profile of cytokines independent of the type of virus (Figure 7). In both viral types, we could observe high levels of IL-6 and IL-8. The rise in IL-6 was expected since it is known that MSCs can secrete various cytokines, such as IL-6, that have been reported to directly affect cell death pathways. The increase in IL-6 and IL-8 in the supernatants suggests that, when infected, GeLo cells might be involved in modulating the local and systemic immune response, which may contribute to resolution of the infection. The low or absent expression of IL-1β in the supernatant of DNA and RNA, respectively, virus-infected GeLo cells may reflect modulation of the immune response, which may be beneficial in some contexts. IL-1β is a key pro-inflammatory cytokine in the early activation of the immune response and induction of inflammation, but its absence or low levels may be beneficial in reducing excessive inflammation. In the case of viral infection, uncontrolled IL-1β production could lead to severe systemic inflammation, increasing the risk of tissue damage and associated complications. Reducing IL-1β could therefore contribute to a more controlled immune response and prevent chronic or harmful inflammatory damage. The higher levels of IL-4 in the supernatant of GeLo cells infected with DNA viruses compared to RNA viruses may be due to differences in the way these two types of viruses activate immune responses. DNA viruses are generally recognised by TLR-9 in endosomes, triggering an immune response that may be less pro-inflammatory in its early phase compared to RNA viruses. In addition, DNA virus infection often triggers a more rapid adaptive response, including activation of T and B lymphocytes, and may promote polarisation. Thus, increased IL- 4 production in cells infected with a DNA virus reflects an immune response that is more focused on activating the humoral response. All these findings could support the use of these cells as treatments in infectious diseases.

To conclude, one of the possible applications of the GeLo cell line may be related to the adaptation of viral vaccine vectors such as MVA (32) or Adenovirus with the purpose of adapting the viral vectors to ovine cells and improving their potential as vaccine candidates in ruminants. One example could be the MVA virus that we have used in previous works to develop vaccines against RVFV and although the results obtained were promising, further optimization should be carried out (44,45). Other applications include the isolation and characterization of viruses and virus-host interactions.

## 5. Conclusions

In this work, we used isolated MSCs from peripheral blood to establish a new ovine cell line to use it as a tool to isolate or produce virus from the same origin as the host. The characterization and the osteogenic and neuronal differentiation carried out confirmed that the GeLo cell line had a mesenchymal stromal origin. In fact, the new established ovine cell line was permissive to all viruses studied in this work. The new sheep cell line could be a useful tool for isolating and characterizing viruses and studying virus-host interactions.

## Supporting information

data figure results

## Acknowledgements

We sincerely thank Dr Javier Castillo from Pirbright Institute for kindly supply the Akabane virus; Dr Borrego from Centro de Investigacion en Sanidad Animal for providing FMDV and sera; Dr Sanchez Seco from Instituto de Salud Carlos III for providing Toscana virus; Dr Nicola Brescia from BIOGUNE for providing Schmallenberg virus; Dr Jimenez Clavero and Dr Llorente for providing the Peste du Petits ruminant virus and West Nile virus; Celia Alonso PhD student for her support.

## Funding

This work was funded by Grupos con Potencial de Crecemento, Xunta de Galicia (ED431B 2023/58 and ED431B 2020/55); and Fundación Pública Gallega de Investigación Biomédica (INIBIC)-Proyectos de Desarrollo y Transferencia 2024 and 2022 and 2024ICT283 “ayudas extraordinarias a la incorporación de científicos titulares” from the Consejo Superior de Investigaciones Científicas (CSIC). We declare no competing interests.

## Contributions

GLA and SD-P designed this study. SM, CSR, SRF, SDP and GLA carried out most of the experiments and data analysis. ECP, AML, AN, JO and LFJ helped with in vitro experiments. GLA and SM prepared the manuscript and the rest of authors helped with paper editing and format revision. The final manuscript was approved by all authors.

Corresponding authors

Correspondence to Gema Lorenzo, Silvia Díaz-Prado or Sandra Moreno

## Data availability

The data that support the findings of this study are available from the corresponding author upon reasonable request.

## Ethics declarations

### Consent for publication

All the authors give their consent for publication in Stem Cell Research & Therapy.

### Ethical approval

All experiments with sheep were performed at Animal Health Research center (CISA-INIA-CSIC) following EC guidelines [Directive 86/609) upon approval by the Ethical Review and Animal Care Committees of INIA and Comunidad de Madrid (authorization decision PRO-EX 192/17).

### AI usage

The authors declare that they have not used artificial intelligence.

### Conflict of interest

The authors declare that they have no conflicts of interest.

## Notes

### Competing Interest Statement

The authors have declared no competing interest.

