## supplementary figure for "Generation and characterization of an ovine cell line derived from peripheral blood and its potential use to study livestock and zoonotic viral infections"

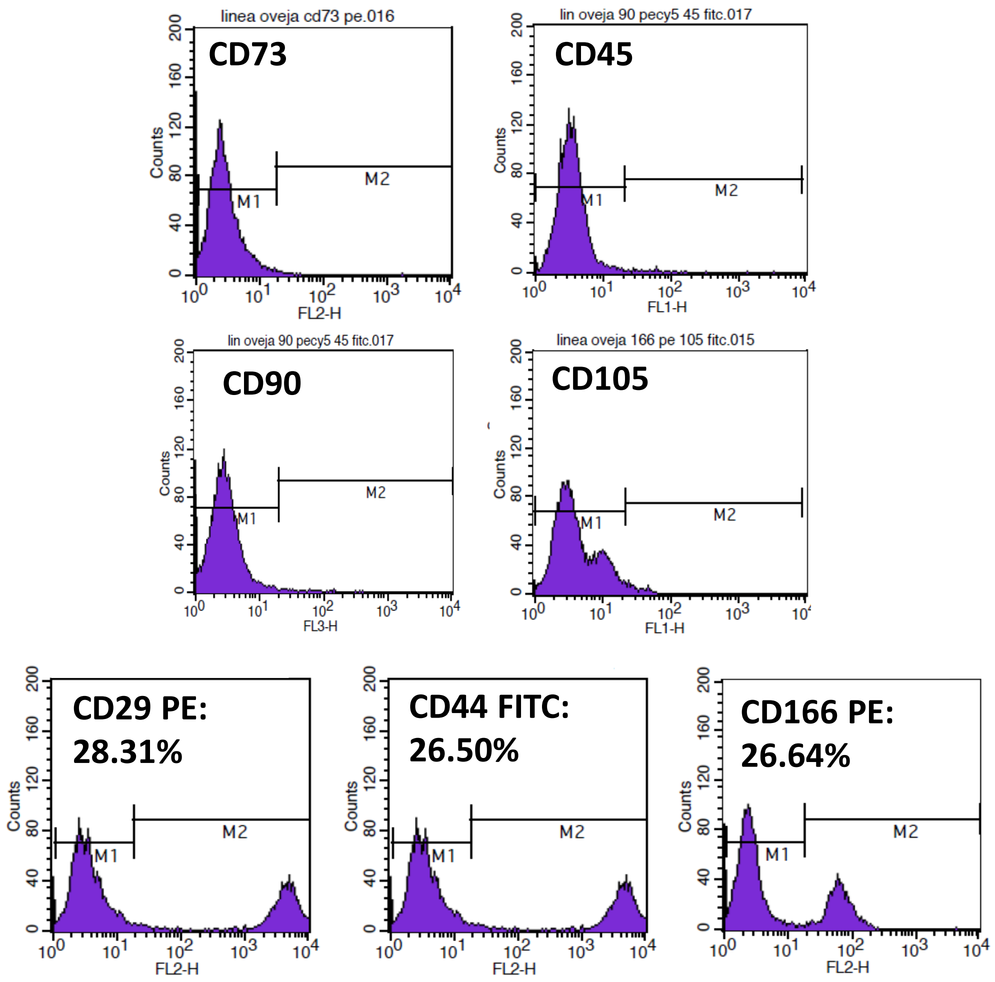


**Supplementary figure 1.** Phenotypic characterization by flow cytometry of GeLo cell line (prior to cloning) for markers characteristic of MSCs (CD29, CD44, CD73, CD90, CD105 and CD166) and hematopoietic cells (CD45) . GeLo cell line was positive for CD29, CD44 and CD166, and negative for CD45, CD73, CD90 and CD105.
