## Supplementary figures and images for "Generation and characterization of an ovine cell line derived from peripheral blood and its potential use to study livestock and zoonotic viral infections"

### supplemental figure 2

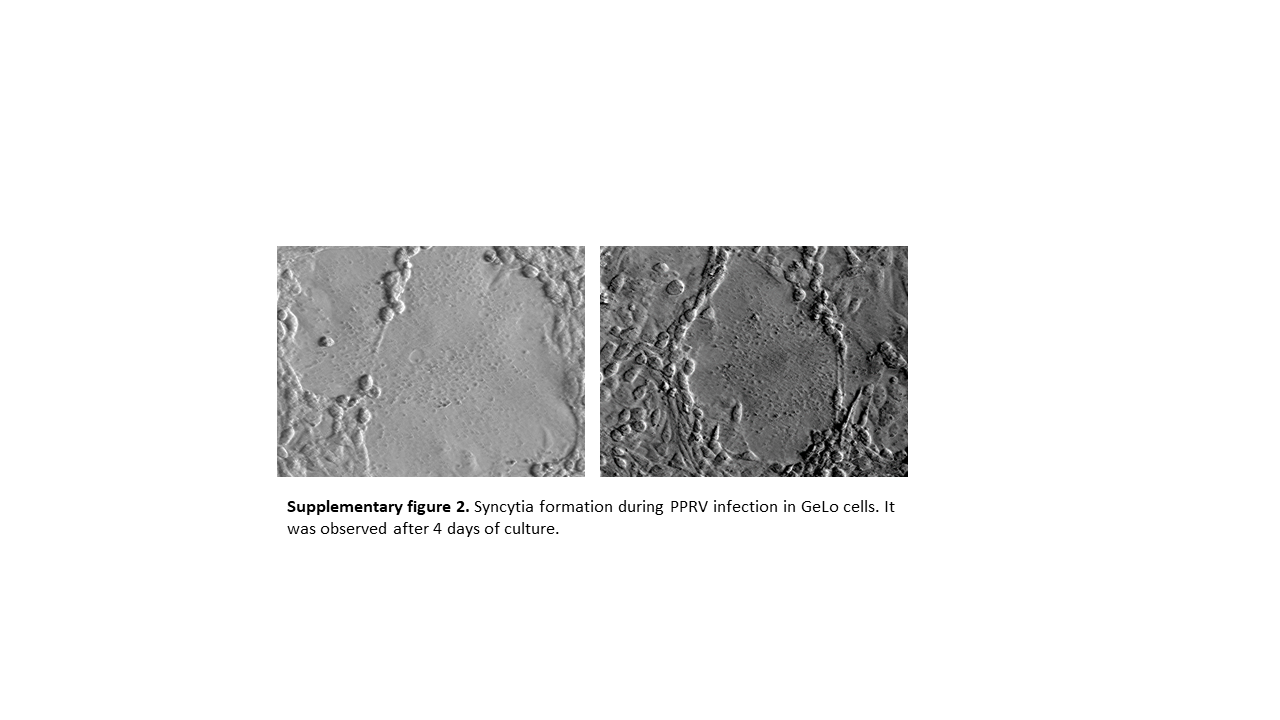
